# Modeling the Electro-chemical Properties of Microbial Opsin ChrimsonR for Application to Optogenetics-based Vision Restoration

**DOI:** 10.1101/417899

**Authors:** Quentin Sabatier, Corentin Joffrois, Grégory Gauvain, Joël Chavas, Didier Pruneau, Serge Picaud, Ryad Benosman

## Abstract

Optogenetic activation of neurons [1] have greatly contributed to our understanding of how neural circuits operate, and holds huge promise in the field of neural prosthetics, particularly in sensory restoration. The discovery of new channelrhodopsins, Chrimson — which is 45 nm more red-shifted than any previously discovered or engineered channelrhodopsin — and its mutant ChrimsonR with faster kinetics [2] made this technology available for medical applications. However, a detailed model that would be able to accurately reproduce the membrane potential dynamics in cells transfected with ChrimsonR under light stimulation is missing. We address this issue by developing the first model for the electrochemical behavior of ChrimsonR that predicts its conductance in response to arbitrary light stimulation. Our model captures ON and OFF dynamics of the protein for stimuli with frequencies up to 100 Hz and their relationship with the brightness, as well as its activation curve, the steady-state amplitude of the response as a function of light intensity. Additionally, we capture a slow adaptation mechanism at a timescale at the order of minutes. Our model holds for light intensities covering the whole dynamic range of the channel (from response onset to saturation) and for timescales in the order of up to several minutes. This model is a new step towards modeling the spiking activity of ChrimsonR-expressing neurons, required for the precise control of information transmission in optogenetics-based Brain-Computer Interfaces, and will inform future applications of ChrimsonR based optogenetics.

## Introduction

This paper studies the activation dynamics of ChrimsonR, a channelrhodopsin which is currently the best candidate for optogenetic-based sensory restoration in human patients, due to its peak spectral sensitivity in the red at 590 nm, which is 45 nm more red-shifted than all previously known channelrhodopsins [3]. However, design of light-stimulation protocols for optogenetics-based vision restoration would greatly benefit from a model of the channel’s behavior, as its dynamics will influence the ChrimsonR mediated temporal dynamics of membrane potential in transfected neurons. To address this issue, here we introduce for the first time the relevant kinetics and the expected behavior of the channel. In particular, we propose a predictive model of the conductance of a population of channels expressed in a given cell in response to arbitrary illumination stimuli spanning the whole dynamic range of the protein.

Upon photostimulation, channelrhodopsins modify their conformation and form a water-filled pore allowing some ions to flow through the channel. The illumination signal therefore drives the conductance of the cell’s membrane, which then induces a current through the membrane, the subsequent modification of the membrane potential and the possible triggering of action potentials. A predictive model of the channelrhodopsin’s conductance provides the mapping between the controlled illumination and the physiologically relevant variable.

The main features of this transformation that we capture are the ON and OFF dynamics (and their relationship to the brightness value of the stimulus); the *activation curve*, the steady-state amplitude of the response as a function of light intensity as well as a slow adaptation mechanism leading to a progressive decrease in the amplitude of the response. These features can be captured using a five-state Markov kinetic model [4]. It is able to predict accurately the response of the protein on long timescales (several minutes) for temporal frequencies lower than approx. 100 Hz and over the whole dynamic range of the protein (between no response at 10^15^ ph.s^−1^.cm^−2^ and saturation around 10^19^ ph.s^−1^.cm^−2^).

The model was primarily build based on the important effort devoted to understanding the chemical reaction underlying the functional behavior of channelrhodopsins. Studies include (i) spectroscopy to determine the relevant conformations of the protein during light excitation, (ii) crystallography which reveals the precise structure of the protein and location of the amino acid residues and retinal chromophore [5, 6], (iii) electrophysiology to study the dynamics of the conductance induced by light stimulation [7–10], (iv) numerical simulations [11–13] and (v) site-directed mutagenesis [14–18] targeting putative key residues to test their influence on experimental observations and infer their role in the photocycle.

Together, this body of work established the outline of the photocycle [19, 20]. Starting from the dark-adapted conformation, and upon photo-induced isomerization of the retinal chromophore, the protein goes through a number of intermediate states finally leading to the conductive state. From this conductive state, the protein either relaxes to the initial dark state or to another closed state, which may or may not be photo-sensitive, and which relaxes slowly (approx. 20 seconds) to the fully dark-adapted state [21].

However, this single-loop photocycle cannot account for a number of observations from electrophysiological (voltage-clamp) experiments. In particular, when the light is switched off, the conductance returns to the baseline according to a linear combination of two exponential terms, with two distinct time constants [11, 12]. This phenomenon implies that the channel must have two distinct conformations which are conductive. This conclusion is consolidated by voltage-clamp experiments at different holding potentials showed that the cation-selectivity of the two open states were different, as shown by the distortion of the conductance responses for different holding potentials [9, 22].

Since then, the prevailing hypothesis has been that the full photocycle consists in two similar half cycles, each one consisting of twin states which are not distinguishable based on spectroscopic analysis [8, 15, 23, 24]. The question remains of where the transitions occur between the two half cycles, and whether they are thermal or light-induced. It was first proposed that the transitions occur between the inactivated states [24]. Recently, it was shown that a light-induced transition existed between the two dark adapted states [25]. Finally, a side reaction was revealed in [23] on the photocycle of the ChR-2 mutant C128T. This side reaction was later added to the two-photocycle model [24]. We believe that this side reaction is responsible for the slow adaptation mechanism we observe.

Our model reduces each half cycle to a simple pair with one closed and one open state. Following conclusions from [25], we set the transitions between the two cycles between the closed dark-adapted states. These transitions are light-induced. Under this form, the model is close to previous modeling work on Channelrhodopsin-2 [11, 12]. We added the side reaction to account for the slow adaptation mechanism. It involves an additional state capturing a fraction of the channels in the conductive state from the second half cycle, and relaxes slowly into the dark-adapted state of the first half cycle.

The numerical parameters of our model were estimated using a least-square procedure between the simulated conductances and recordings from patch-clamp experiments performed on ChrimsonR-expressing HEK293 cells at physiological temperature. The usage of HEK293 cells is justfied by previous electrophysiological experiments in different cell types including *Xenopus* oocytes, human embryonic kidney (HEK) cells, baby hamster kidney (BHK) cells, Henrietta Lacks (HeLa) cells, cultured neurons, etc… which have shown that the ChrimsonR properties to be mostly insensitive to the host system used [10, 12]. This model is an important step towards modeling the spiking activity of ChrimsonR-expressing neurons, required for the precise control of information transmission in optogenetics-based Brain-Computer Interfaces, and will inform future ChrimsonR based optogenetic applications.

## Methods

### HEK 293T cell culture, transfection

HEK 293T cells were maintained between 10% and 70% confluence in DMEM medium (Invitrogen, Waltham, USA) supplemented with 10% FBS (Invitrogen), 1% penicillin/streptomycin (Invitrogen). For recording, cells were plated at 50,000 cells per well in 24-well plates that contained round glass coverslips (12 mm) coated with polylysine (2 µg.cm^−2^, Sigma Aldrich) and laminin (1 µg.cm^−2^, Sigma Aldrich). Adherent cells were transfected approximately 24h post-plating with JetPrime (Polyplus Transfection) and recorded via whole-cell patch clamp between 24 and 72h post-transfection. 0.5 µg of DNA was delivered per well. In addition, all-trans retinal (ATR, 10 µM) was supplemented to the culture medium for 1h before patch-clamp experiments.

The opsin ChrimsonR was expressed into the HEK cells associated with a fluorescent protein tdTomato fused to its C-terminal end. Transfection was performed through the mutant capsid AV2-7m8 [26]. This construct was chosen for its efficiency in transfecting retinal ganglion cells through intravitreal injection.

### Electrophysiolgy

Whole-cell patch-clamp recordings were performed in isolated HEK 293T cells to avoid space clamp issues. All recordings were performed using an Axopatch 200B amplifier and Digidata 1440 digitizer (Molecular Devices) at room temperature. Steady access resistance were inferior to 40 MΩ. Typical membrane resistance was between 200 MΩ and 1 GΩ, and pipette resistance was between 5 and 8 MΩ. Cells were perfused with Ames medium (Sigma-Aldrich, St Louis, MO; A1420) bubbled with 95% O_2_ and 5% CO_2_ at 37 °C at a rate of 1-2 ml.min^−1^ during experiments. Intracellular solution was composed of (in mM) 115 K-gluconate, 10 KCl, 0.5 CaCl_2_, 1 MgCl_2_, 1.5 EGTA, 10 HEPES, 4 ATP-Na_2_, pH 7.3 (KOH adjusted).

### Illumination

Photostimulation of patch-clamped cells was conducted with a 595 nm LED (M595L3, Thorlabs, half bandwidth of 75 nm). Irradiance was measured, for different LED voltages, at the level of the coverslip sample using a power meter composed of a digital optical power and energy meter (Thorlabs, PM100D) and a photodiode power sensor (Thorlabs, S120C). Resulting power measurements were converted in ph.s^−1^.cm^−2^ using known illuminated surface (1.21 mm^2^) and peak wavelength for the LED (595 nm). These measurements were confirmed using a calibrated spectrophotometer (USB2000+, Ocean Optics, in-house calibration).

### Illumination protocols

The building block of our stimulation protocol is a series of ten 200 ms square pulses repeated at a frequency of 0.5 Hz. These building blocks were then assembled in several ways with different light levels. The duration between two subsequent blocks was not set *a priori* (as it required replacing manually an optical filter between the light source and the recorded cell) but the actual value was recorded and usually lasted about ten seconds.

Light intensity of stimulation blocks covered the whole range of responses from the protein, from virtually no response at 3×10^15^ ph.s^−1^.cm^−2^ to saturation in amplitude at 10^19^ ph.s^−1^.cm^−2^.

### Data analysis and simulation

#### Markov kinetic models

The behavior of the protein ChrimsonR is modeled using a Markov kinetic model. In this model, a number of *states* represent the different conformations that the protein can take. For each pair of states, there can be a transition between these two states if the protein can switch from the first state to the other without going through any other stable conformation. It is worth noting that transitions are not necessarily reversible, i.e. if the transition from state *i* to state *j* exists, the transition from *j* to *i* does not always exist. Additionally, a time constant is associated with each transition. For a pair of states (*i, j*), the rate *λ*_*i, j*_ — the inverse of the time constant *T*_*i, j*_ — quantifies the probability that the protein jumps from state *i* to state *j* per unit of time. If there is no transition from state *i* to state *j*, this transition has a zero transition rate, or equivalently an infinite time constant.

For a given model with *N* states, and a transition matrix 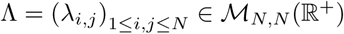, and denoting *p*_*i*_(*t*) the probability that the protein is in state *i* at time *t*, the evolution of the system is given by *P* (*t*) = (*p*_*i*_(*t*))_1 ≤ *i* ≤ *N*_ and follows the system:

*∀i ∈* ⟦1, *N* ⟧

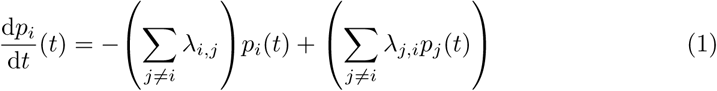

In the rest of the paper, we will consider two types of transitions, thermal and photo-induced. Thermal reactions involve absorption or evolution of heat. They take place also in the absence of light. Temperature has a significant effect on the rate of a thermochemical reaction. On the other hand, photochemical reactions involve absorption of light. The presence of light is the primary requisite for the reaction to take place and the rate of a photochemical reaction depends on the intensity of the light. Temperature has very little effect on the rate of a photochemical reaction.

Mathematically, for a given temperature, a thermal reaction is represented by a constant rate, while the rate of the photochemical transition varies linearly with light intensity. Figure (1, a) shows an example of photocycle combining the conclusions from several spectroscopic studies [24, 25] on different channelrhodopsins. The photocycle consists in two separate cycles, each involving a conductive state (P520 and P520’), with possible transitions from one half-cycle to the other and a side reaction (states P380 and P353) with a transition to either of the two half-cycles.

**Fig 1.**
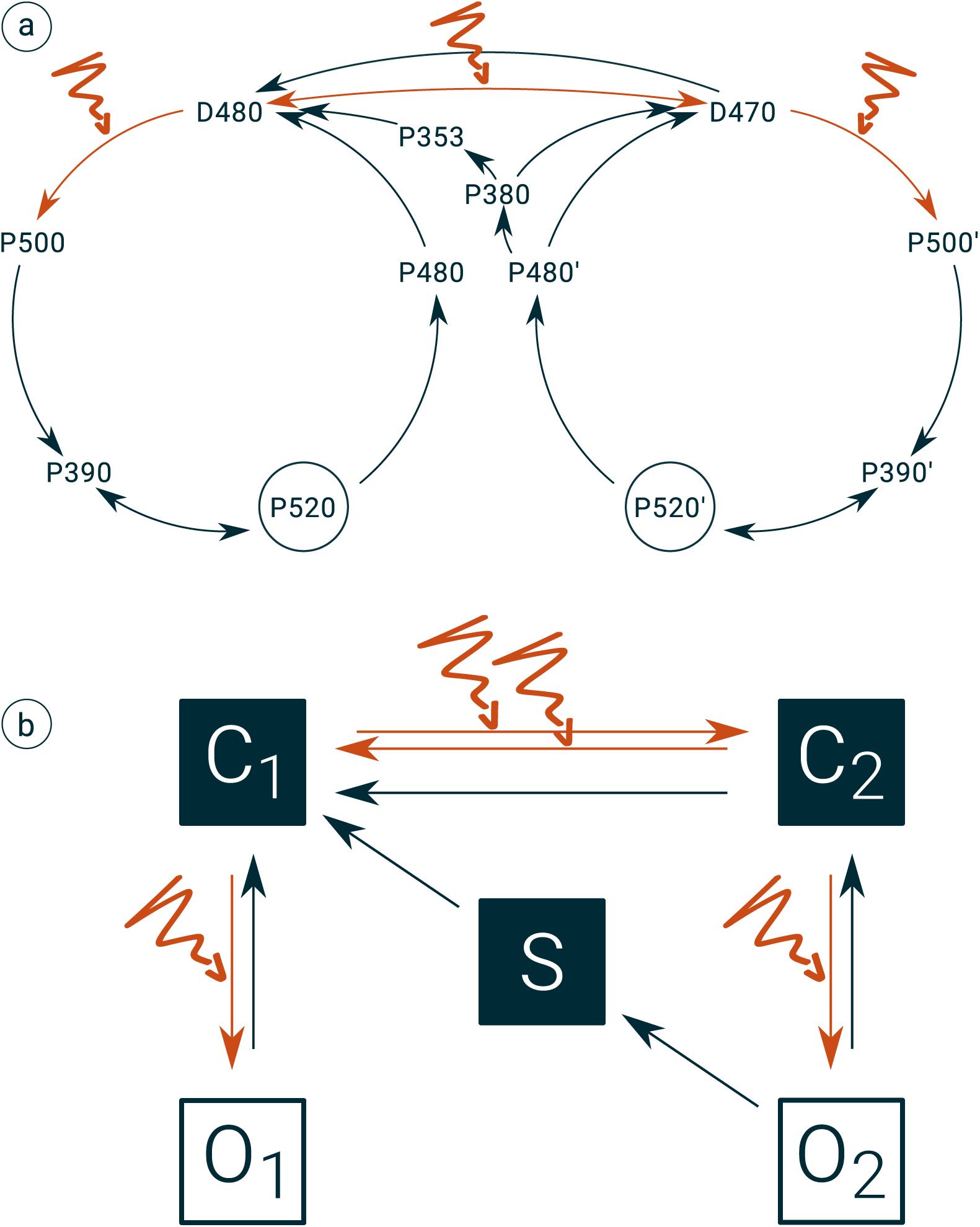
The photocycle (a), such as provided by previous studies on other channelrhodopsins, and the simplified model (b) we used to approximate the electrochemical behavior of ChrimsonR.

Fig. (1, b) shows the Markov chain we used to model the behavior of ChrimsonR. It is a simplified version of the more realistic photocycle introduced above. The two half-cycles are represented by C_1_⇄O_1_ and C_2_⇄O_2_. C_1_ and C_2_ represent the ground states D480 and D470 respectively. The intermediate states P500, P500’, P390 and P390’, which are fast intermediates, have been suppressed. States P520 and P480 have been merged into O_1_, P520’ and P480’ into O_2_. The side reaction is represented by the single state S, and the transition from S back to half-cycle 2 has been omitted because it could not be resolved based on our data.

#### Fitting linear combinations of exponential functions

A general result on continuous-time Markov chains [4] with time-independent transition rates is that, if the Markov chain has a number *N* of states, then the probability *p*_*i*_(*t*) that the chain is in state *i ∈* ⟦1, *N* ⟧ at time *t* can be expressed as:

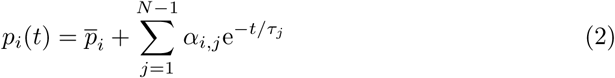

where 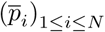 are the stable probabilities for each state, i.e. the probabilities which are invariant by multiplication by the transition matrix. (τ_*j*_)_1 ≤ *j* ≤ *N* − 1_ is a set of *N* − 1 time constants which are shared by the *N* states, and (α_*i, j*_)_1 ≤ *i* ≤ *N*, 1 ≤ *j* ≤ *N* − 1_ are the weights of the linear combination, and depend on the initial conditions. In our case, the transition rates are constant over a time period if and only if light intensity is constant over this period.

Additionally, we do not observe the probabilities directly, but rather a linear combination of the total number of channels which are in each conductive state (P520 and P520’), the weights being the conductances of each state. However, since the number of channels expressed in each patch-clamped cell is constant over time, the fraction of channels in a state *i* is a good approximation of the probability *p*_*i*_. Since the *N* states share the same *N* − 1 time constants, the linear combination only affects the coefficients and thus does not prevent us from estimating the time constants.

Fig. (2) shows that three terms describe well the time course of a single pulse during the light pulse, while two terms are enough to describe the dynamics of the decay when light is turned off. This double-exponential dark decay is reported for all channelrhodopsins in the literature. It is, in fact, the main argument against the single photocycle model, which would be the natural conclusion based on spectroscopic studies only.

**Fig 2.**
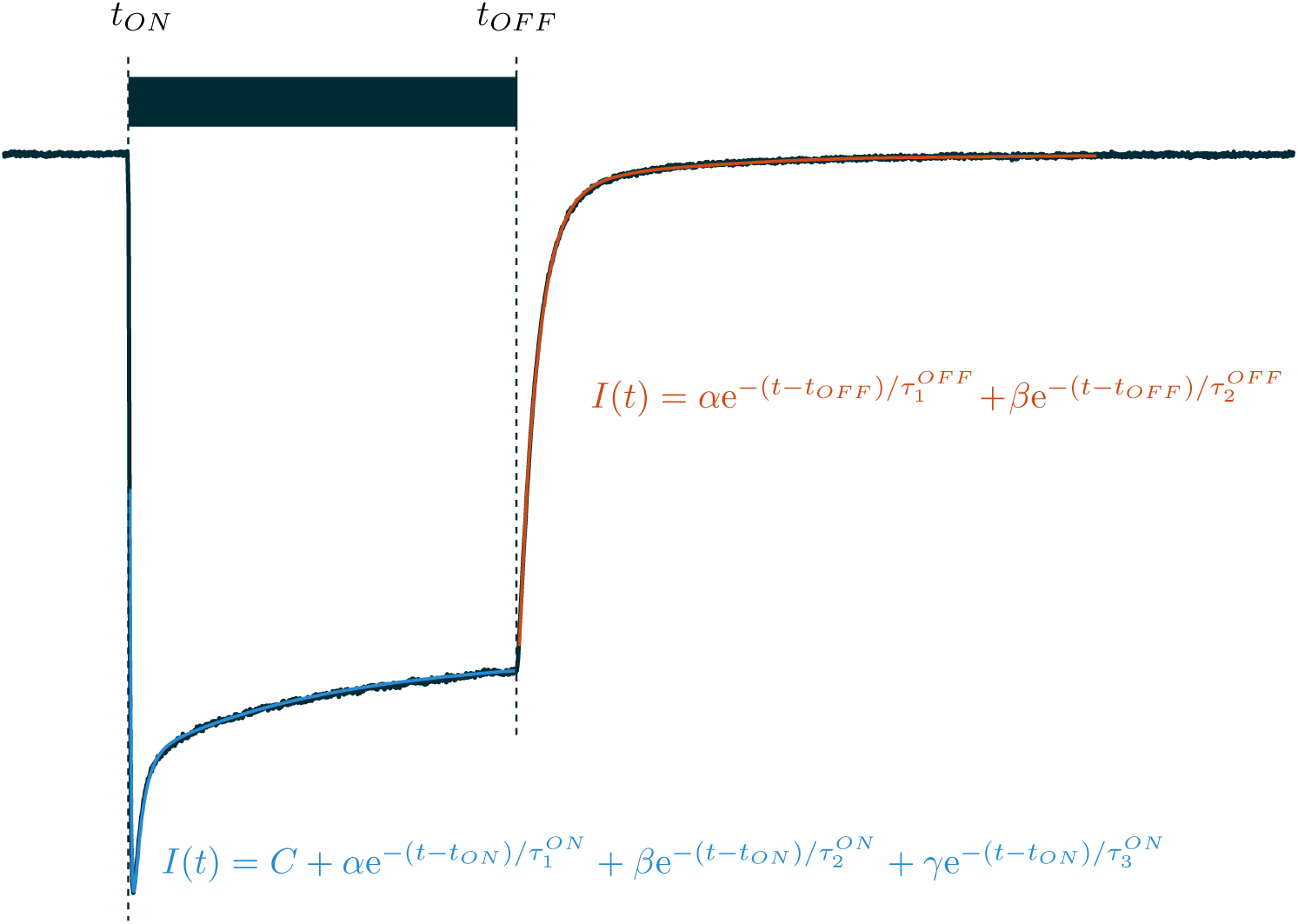
Estimating the On and Off dynamics. Linear combinations of exponential functions were fitted to the portions of the curves where light intensity is constant. The number of terms was fixed based on the shape of the responses. For on dynamics (non-zero light intensity) a constant term is required in order to represent the steady-state level. Given that the steady-state level is known to be zero in the absence of light, the constant term was removed for the analysis of the off dynamics.

**Fig 3.**
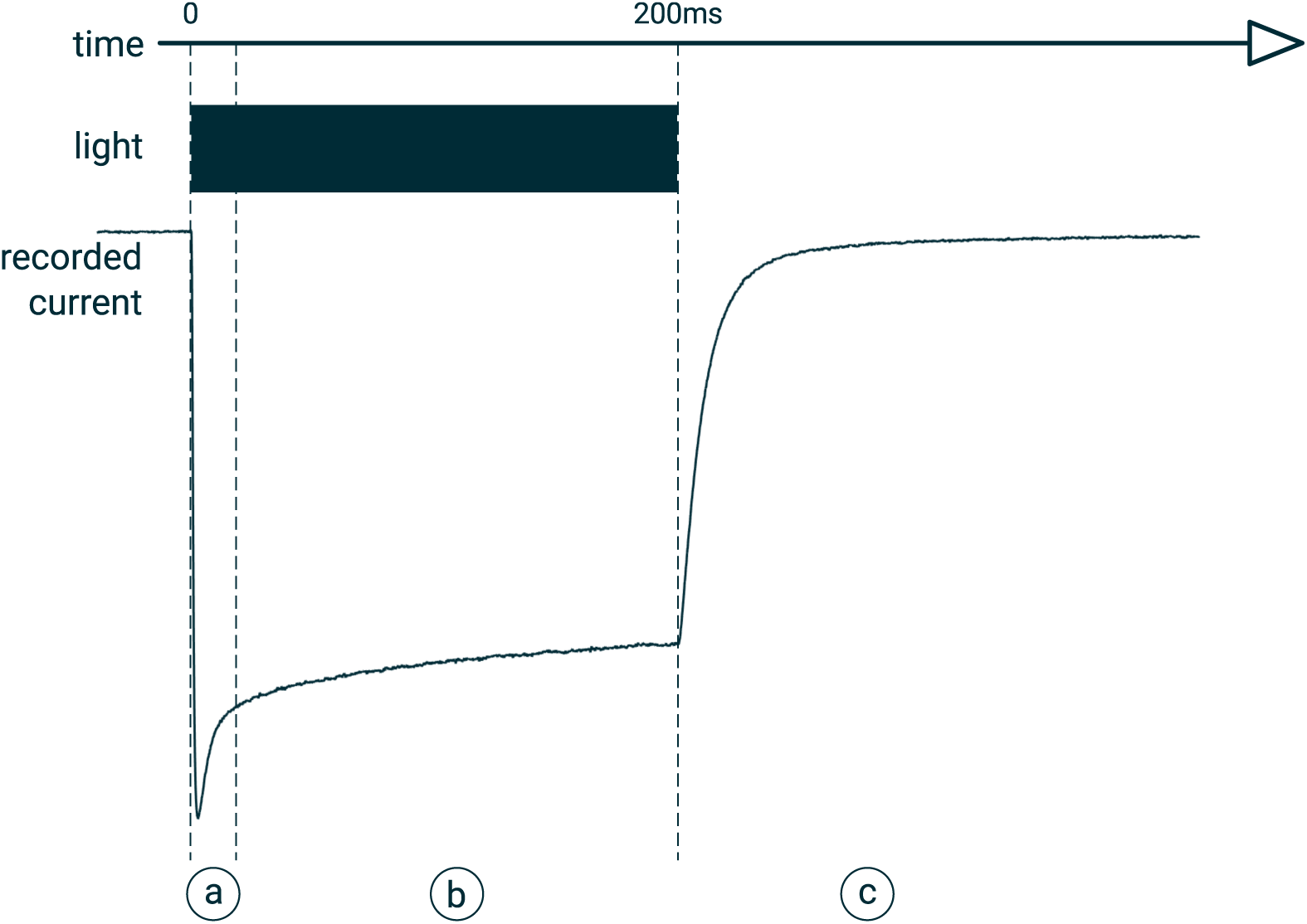
The generic shape of the response to a square pulse of light: (a) a fast response, which depends on the stimulation history and lasts from 10 ms to 40 ms, is followed by (b) a slow decay towards a non-zero equilibrium value. (c) When light is turned off, the recorded current converges towards zero with a double-exponential decay.

**Fig 4.**
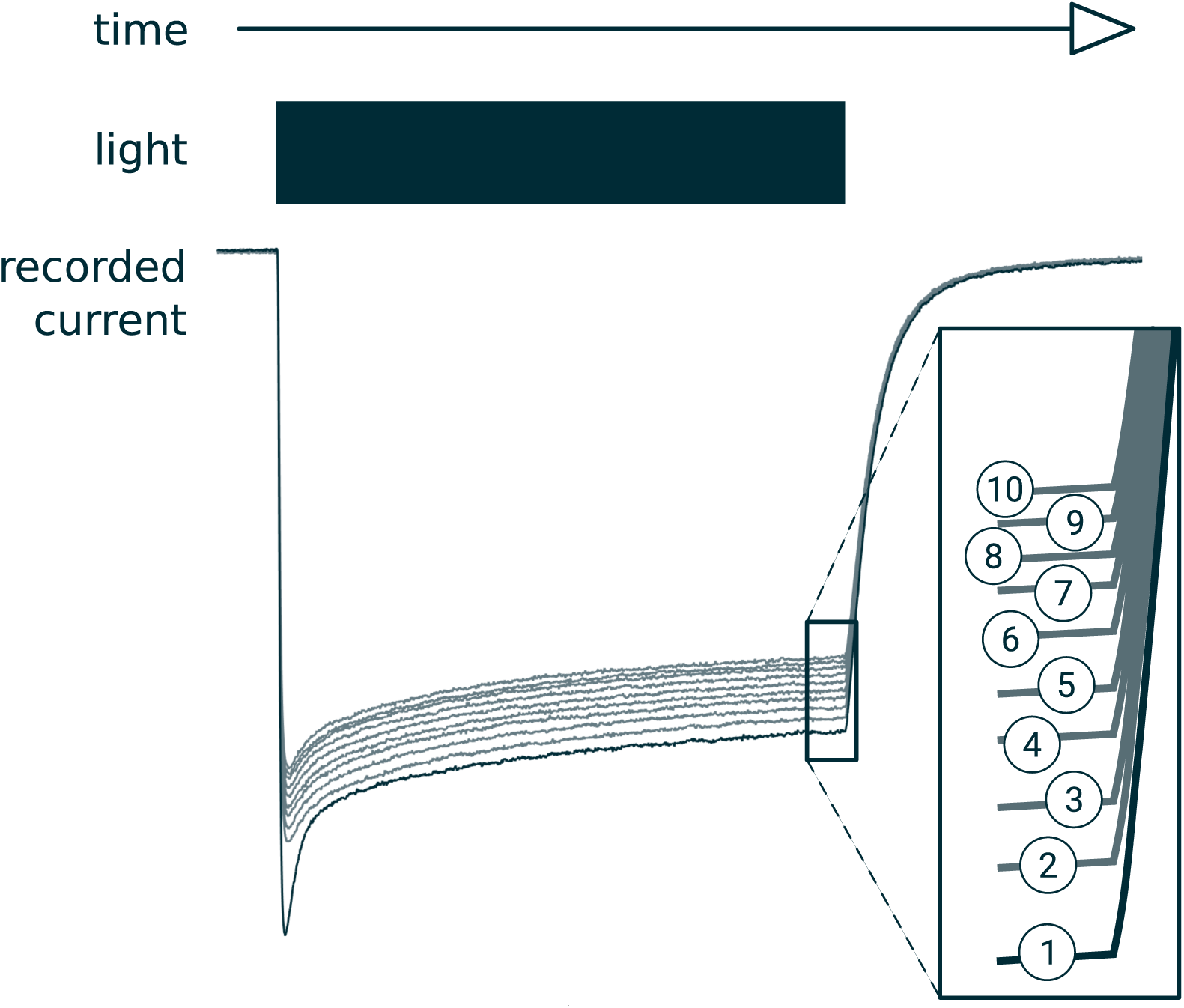
Response to a repeated stimulus (200 ms light pulses at a given intensity —*I* = 1.17×10^19^ ph.s^−1^.cm^−2^ — one pulse every 2 s): the ten responses are synchronized with respect to the stimulus and stacked. The order of the sweeps is given in the box on the right. Except the first pulse, the nine others are only scaled versions of each other.

#### Fitting the parameters of the Markov kinetic model

The parameters of our model were separated into different groups and fitted sequentially.

First, the two off time constants (O_1_ → C_1_ and O_2_ → C_2_) were fixed using the numerical results of the analysis on the linear combinations of exponential functions. In this configuration where the two open states are not connected, the values of the two transitions must be the two time constants describing the off dynamics. Analyzes with the two possibilities showed that the fastest time constant must be associated with the stable cycle C_1_⇄O_1_.

Then the other time constants were estimated through a least-square minimization procedure between the recorded and simulated conductances. The parameters of the error function were the time constants of the transitions, except the transitions O_1_ → C_1_ and O_2_ → C_2_, fixed in the previous step as well as the thermal transition C_2_ → C_1_ which is too slow to be estimated using experiments implemented at this timescale, and which was arbitrarily fixed to a very small value (approx. 30 minutes) so it did not affect the dynamics of the system.

#### Medium-term and long-term activation curves

The side reaction involving the slow state (states P380 and P353 in Fig. (1, a) or S in (1, b)) makes it tricky to estimate the activation curve (steady-state conductance level as a function of the light intensity) of the protein. In fact, it is not trivial to disentangle the influence of the current light intensity level from the history of stimulation on the amplitude of the response. Our estimation of the activation curve relies on the prior fitting procedure of the five-state model to the data. The activation curve can then be recovered through the model, as the steady-state level of the response of the model to the whole range of light intensities.

Here, we present the activation curves at two different timescales. The long-term timescale (several minutes) is enough for the whole chain to converge, including the transitions through the slow state. But we also include a medium-term activation curve, which represents the state of the Markov chain where all transitions but the slow ones have converged. This medium-term equilibrium is obtained by arbitrarily setting the transition rate from O_2_ to S to zero.

The analysis of the time constants describing the response of ChrimsonR according to the five-state model allows us to estimate the time constant of the transition from the medium-term to the long-term equilibria.

## Results

### Main observations

In response to a single 200 ms light pulse, the typical response that we observe has three main features: (i) a fast response occurring in the first 10 ms to 40 ms from pulse onset which can either be an overshoot or an undershoot depending on several factors detailed below, followed by (ii) a slow decay with a time constant of the order of 100 ms (see) towards a non-zero equilibrium value which is not reached within the 200 ms, and (iii) when light is switched off, a decay towards zero characterized by the sum of two exponential terms.

Additionally, when the pulse is repeated (every 2 s in our protocols) at a high intensity (*I*≥10^17^ ph.s^−1^.cm^−2^) the response for a given pulse is generally of the same shape as the previous one but with a slightly smaller amplitude.

Our more valuable observations are derived from clean recordings of the same cell over several minutes, during which the building blocks of the protocols (series of ten pulses) were not arranged in a monotonic order, i.e. block intensity goes up and down (Fig. 10, 12 and 13). In this situation, the shape of the fast part of the response (occurring in the first 10 ms to 40 ms) depends substantially on the light intensity of the previous pulse. Specifically, for a given pulse with a given light intensity, the higher the light intensity during the previous pulse, the lower the fast part of the response (Fig. 5, a). It is our understanding, even if we haven’t tested this hypothesis explicitly, that the time interval between the two pulses has little influence on the fast part of the response. In contrast, the slow part of the response is unaffected by the previous stimulation patterns.

**Fig 5.**
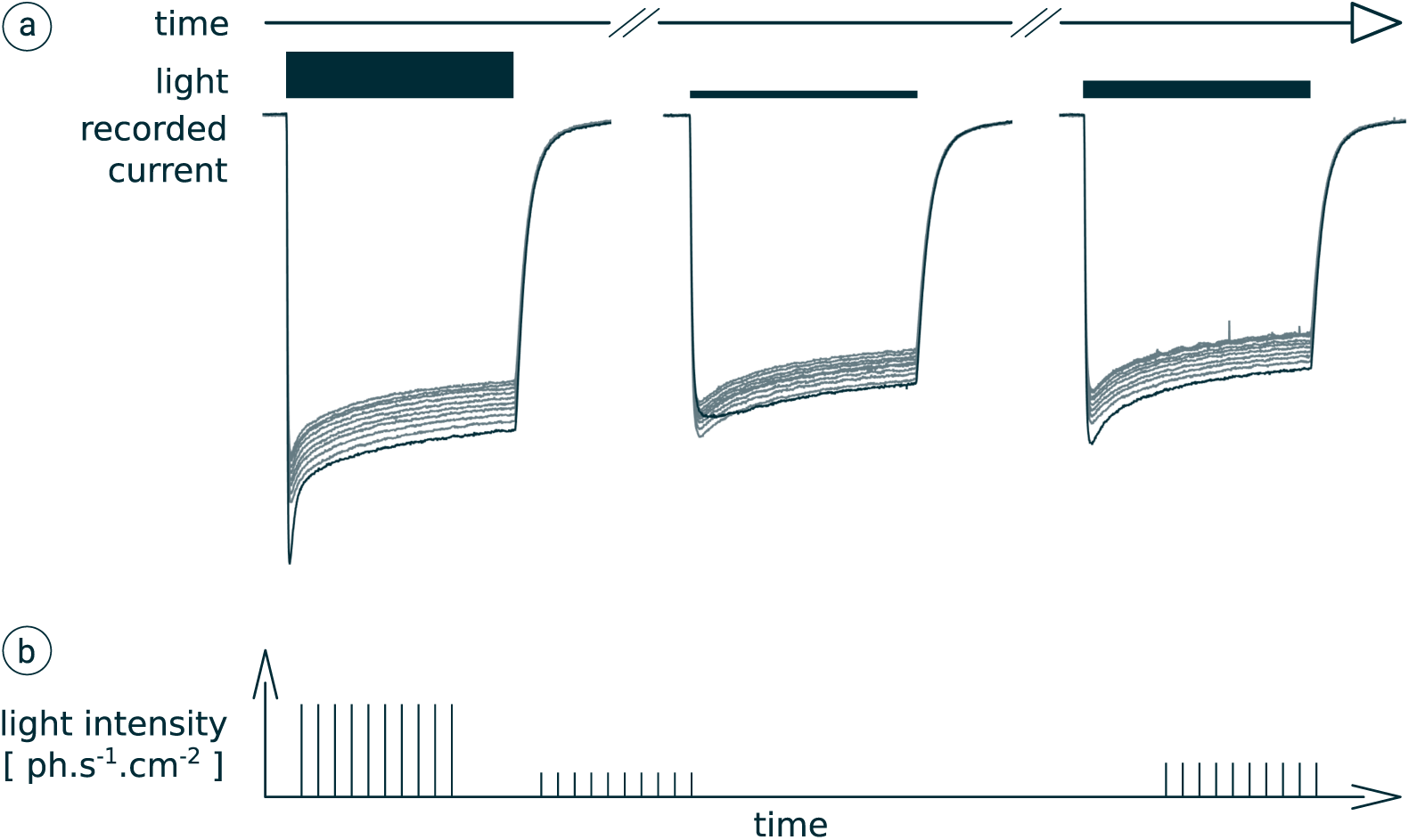
Responses to series of ten 200 ms pulses. (a) Stacked responses: the response to the first pulse is darker. (b) Stimulus description: it consists in three series of ten pulses. In each series, light intensity is constant and one pulse is emitted every 2 s. Light intensities are respectively 1.17 × 10^19^, 3.12 × 10^18^ and 4.36 × 10^18^ ph.s^−1^.cm^−2^.

### Time constants

#### On kinetics

At low light levels (*I* ≤ 5×10^17^ ph.s^−1^.cm^−2^), the time course of the recorded current during an interval when the light is on at a constant intensity is well approximated by the linear combination of (i) a constant term representing the limit current and (ii) two exponential terms with distinct time constants. At higher light levels, an additional exponential term, with a third time constant, is necessary to describe accurately the response to a single light pulse. Fig. (6, b) shows the estimated time constants as a function of light intensity. The fastest time constant (often referred to as τ_*ON*_ in the literature, black curve) — which represents the timescale at which the protein can be activated — is well estimated. It decreases with light intensity, i.e. the higher the intensity, the faster the protein is activated. The estimation of the second time constant is less precise (Fig. (6, b), blue curve). The estimation of the third time constant (orange curve) is very erratic, as shown by the very large confidence intervals on most of the recorded cells on Fig. (6, a). However, the analyzes run on the cleanest data sets suggest that a time constant between 100 ms and 200 ms, relatively independent of the stimulation intensity is a good initial guess.

**Fig 6.**
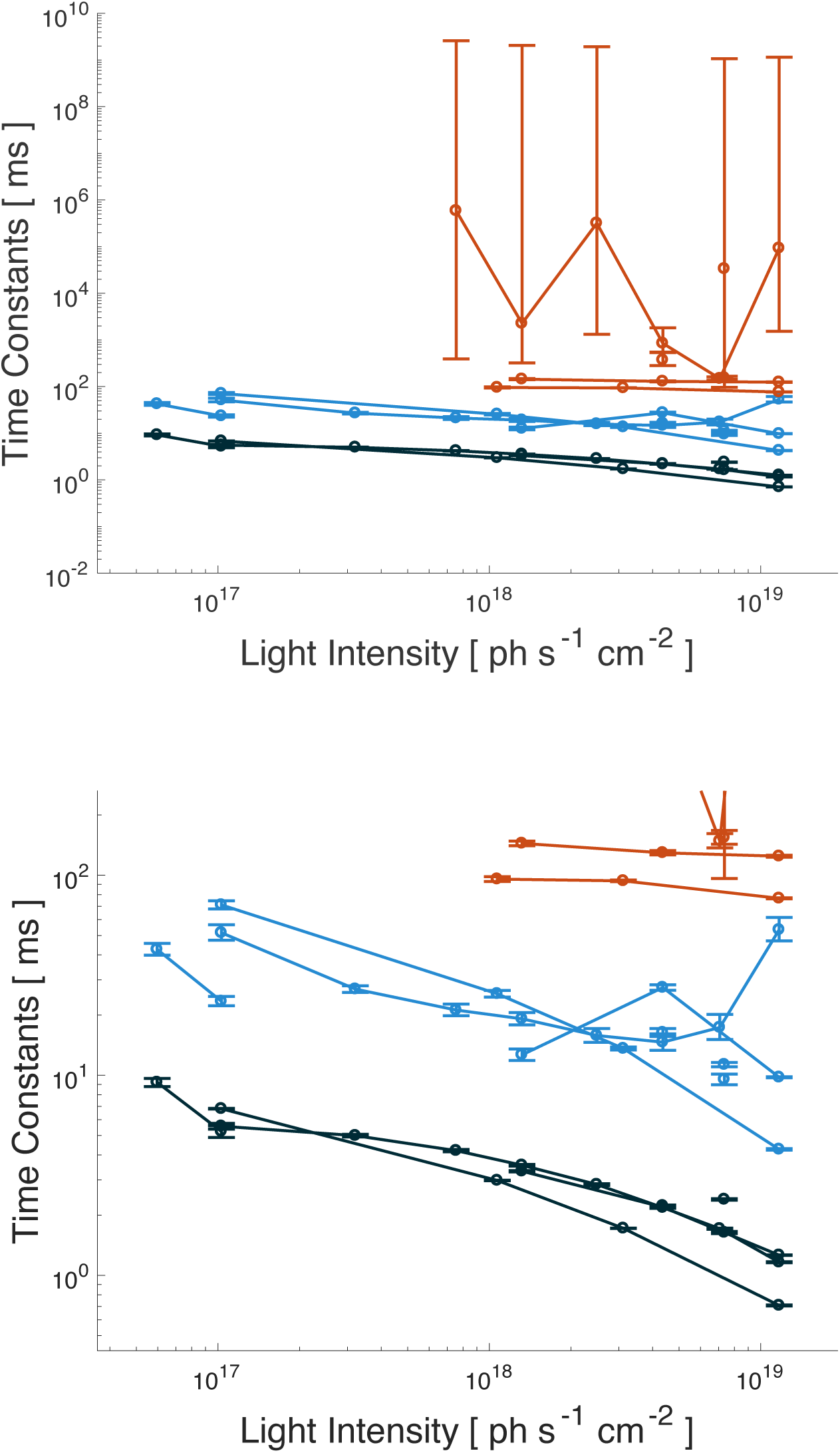
On kinetics analysis of experimental data: linear combinations of exponential functions have been fitted on the recorded responses to light pulses. Top: when available, all three time constants are shown. The precision on the slowest time constant is usually very poor. Bottom: same curve zoomed on the two fastest time constants. The circles show the estimated values for each time constant, and the error bars show the 95% confidence intervals. The data points that are linked together belong to the same cell.

#### Off kinetics

In agreement with what was described in the literature, our data show that the dynamics of the observed current when the light is turned off is very well approximated by a linear combination of two exponential terms. Fig. (7) shows the numerical results obtained separately on each recorded cell.

**Fig 7.**
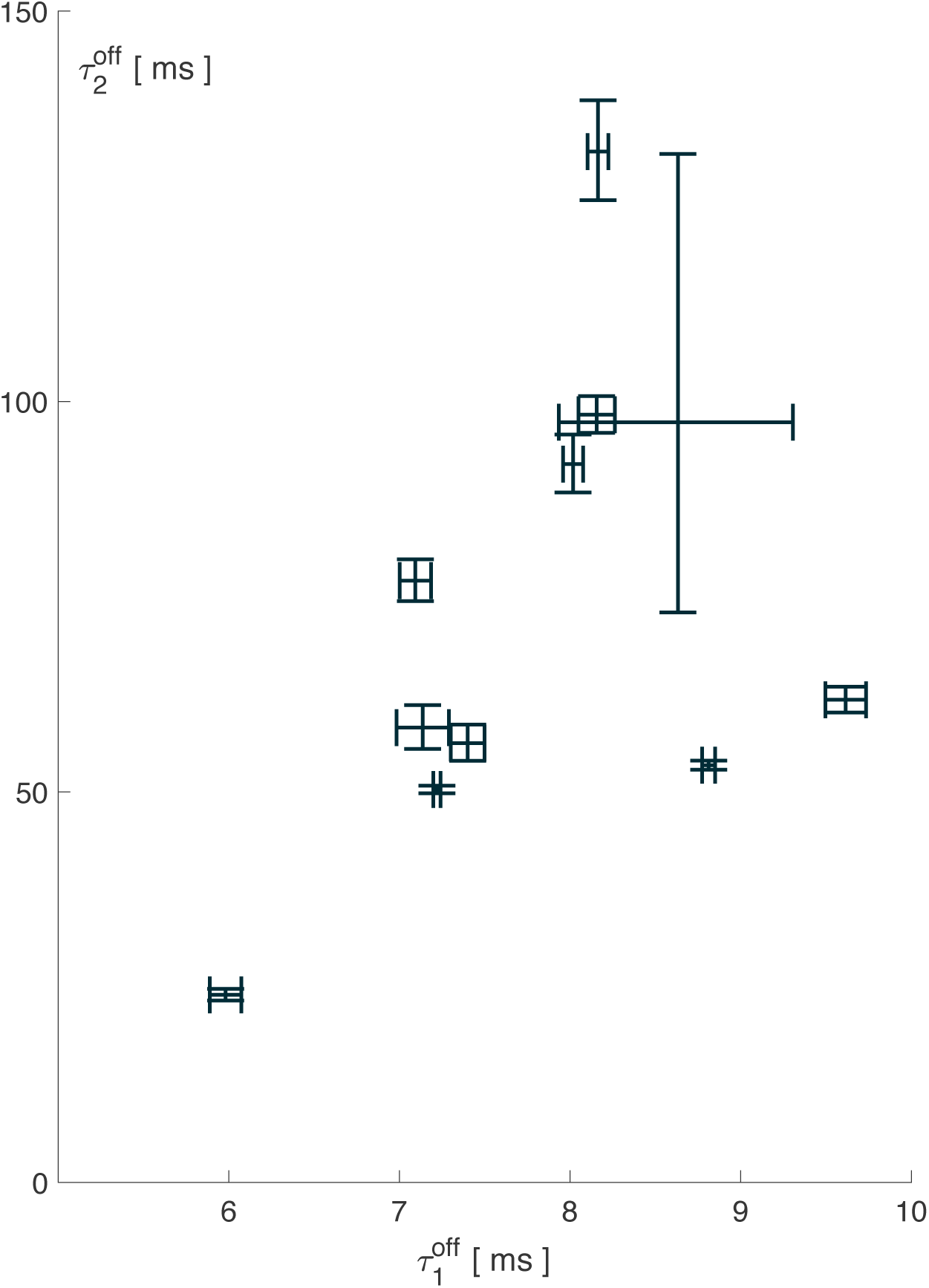
Numerical results for the estimation of the time constants governing the off dynamics of ChrimsonR-tdTomato. For each cell, the estimations of the fast (abscissa) and slow (ordinate) time constants are shown along with the 95% confidence intervals. It is worth noting that intra-cell variability is generally much lower than the inter-cell variability.

### Slow adaptation process

As stated in the *Main observations* section, our data show that the behavior of ChrimsonR involves a slow adaptation process such that the overall amplitude of the response to a series of identical pulses decreases with each pulse. This feature has not been previously reported in electrophysiological experiments, but we think that this phenomenon can be compared with the side reaction involving states P380 and P353 of the photocycle of the channelrhodopsin C128T mutant described in particular in [23, 24].

The timescale at which this adaptation occurs is too long to be observed on single 200 ms pulses. Therefore, the analysis of the on-dynamics (section) cannot bring any valuable information to estimate the time constant of this phenomenon.

Instead, we used the Markov kinetic model involving a slow state representing the side reaction and we fitted the model on the responses of the same cell to stimulation protocols consisting of repeated series of ten pulses. Since the protocols span over several minutes, they allow the estimation of this adaptation process, especially the recovery time constant. Fig. (15) shows side by side the time constants such as estimated directly on single pulses and theoretically using the model after parameter estimation. This second method reveals a fourth time constant, much longer than the three others, and characterizing the slow adaptation process.

### Simplified photocycles

#### Global approximation

The model aims at capturing parts (b) and (c) of the typical response shown in Fig. 3, the overall amplitude of the response including the slow adaption with a time constant of the order of 10^2^ ms and the double exponential decay.

As stated in the *Methods* section, some parameters were fitted individually for each cell, like the time constants from the open states back to the closed states, other parameters were fitted on the most relevant data sets and then used for all parameter sets. Numerical values are provided in tables (1), (2) and (3).

**Table 1.** Estimated conductances for the six different cells (in nS)

| Cell | $g_1$ | $g_2$ |
| --- | --- | --- |
| 1 | 17 | 3.3 |
| 2 | 3.92 | 0.44 |
| 3 | 4.85 | 2.01 |
| 4 | 3.19 | 0.39 |
| 5 | 15.4 | 2.88 |
| 6 | 8.07 | 2.84 |

**Table 2.** Estimated photochemical transition rates (in ms^−1^ / ph.s^−1^.cm^−2^)

| Cell | $C_1 \rightarrow O_1$ | $C_2 \rightarrow O_2$ | $C_1 \rightarrow C_2$ | $C_2 \rightarrow C_1$ |
| --- | --- | --- | --- | --- |
| 1 | $1.61 \times 10^{-19}$ | $3.03 \times 10^{-20}$ | $1.16 \times 10^{-20}$ | $5.99 \times 10^{-20}$ |
| 2 | $1.67 \times 10^{-19}$ | $3.89 \times 10^{-20}$ | $1.28 \times 10^{-20}$ | $6.17 \times 10^{-20}$ |
| 3 | $1.15 \times 10^{-19}$ | $5.84 \times 10^{-20}$ | $2.96 \times 10^{-21}$ | $4.07 \times 10^{-20}$ |
| 4 | $4.60 \times 10^{-19}$ | $1.23 \times 10^{-19}$ | $5.13 \times 10^{-20}$ | $1.46 \times 10^{-19}$ |
| 5 | $1.10 \times 10^{-19}$ | $7.20 \times 10^{-20}$ | $1.94 \times 10^{-21}$ | $1.44 \times 10^{-20}$ |
| 6 | $1.60 \times 10^{-19}$ | $3.87 \times 10^{-20}$ | $4.93 \times 10^{-20}$ | $1.75 \times 10^{-19}$ |

**Table 3.** Estimated thermal transition rates (in ms^−1^)

| Cell | $O_1 \rightarrow C_1$ | $O_2 \rightarrow C_2$ | $O_2 \rightarrow S$ | $C_2 \rightarrow C_1$ | $S \rightarrow C_1$ |
| --- | --- | --- | --- | --- | --- |
| 1 | 0.14 | $1.14 \times 10^{-2}$ | $48.1 \times 10^{-5}$ | $10^{-7}$ | $5.91 \times 10^{-6}$ |
| 2 | 0.12 | $1.78 \times 10^{-2}$ | $7.89 \times 10^{-5}$ | $10^{-7}$ | $3 \times 10^{-6}$ |
| 3 | 0.13 | $1.78 \times 10^{-2}$ | $8.56 \times 10^{-5}$ | $10^{-7}$ | $3 \times 10^{-6}$ |
| 4 | 0.10 | $1.40 \times 10^{-2}$ | $11.5 \times 10^{-5}$ | $10^{-7}$ | $2.49 \times 10^{-6}$ |
| 5 | 0.12 | $1.78 \times 10^{-2}$ | $8.79 \times 10^{-5}$ | $10^{-7}$ | $3 \times 10^{-6}$ |
| 6 | 0.10 | $1.37 \times 10^{-2}$ | $19.2 \times 10^{-5}$ | $10^{-7}$ | $1.5 \times 10^{-6}$ |

Fig. (8), (9), (10), (11), (12) and (13) display the comparison between the data and the simulations for six different cells. They show that the main features are in fact captured by the five-state model.

**Fig 8.**
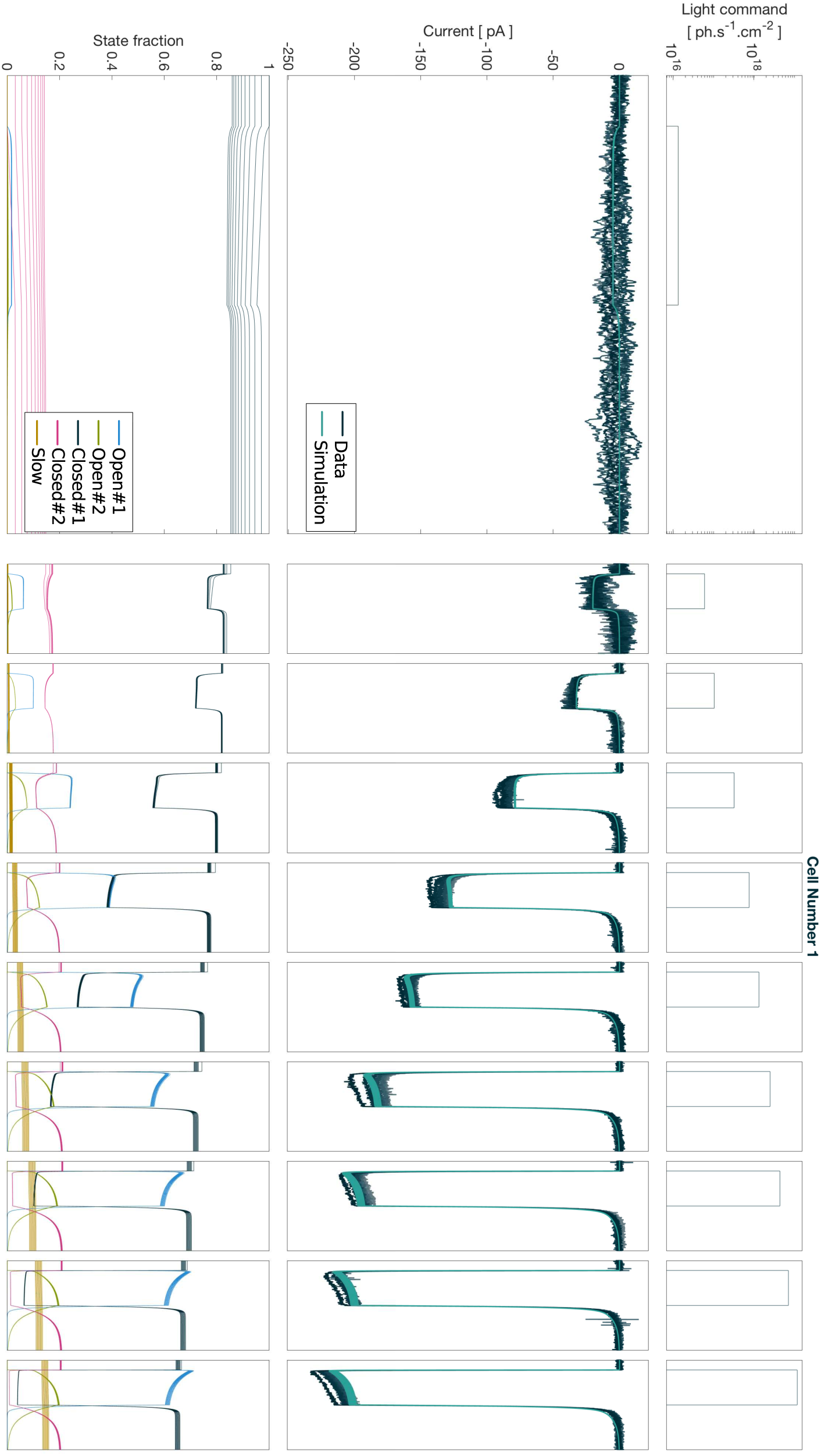
Data and simulation comparison Cell 1.

**Fig 9.**
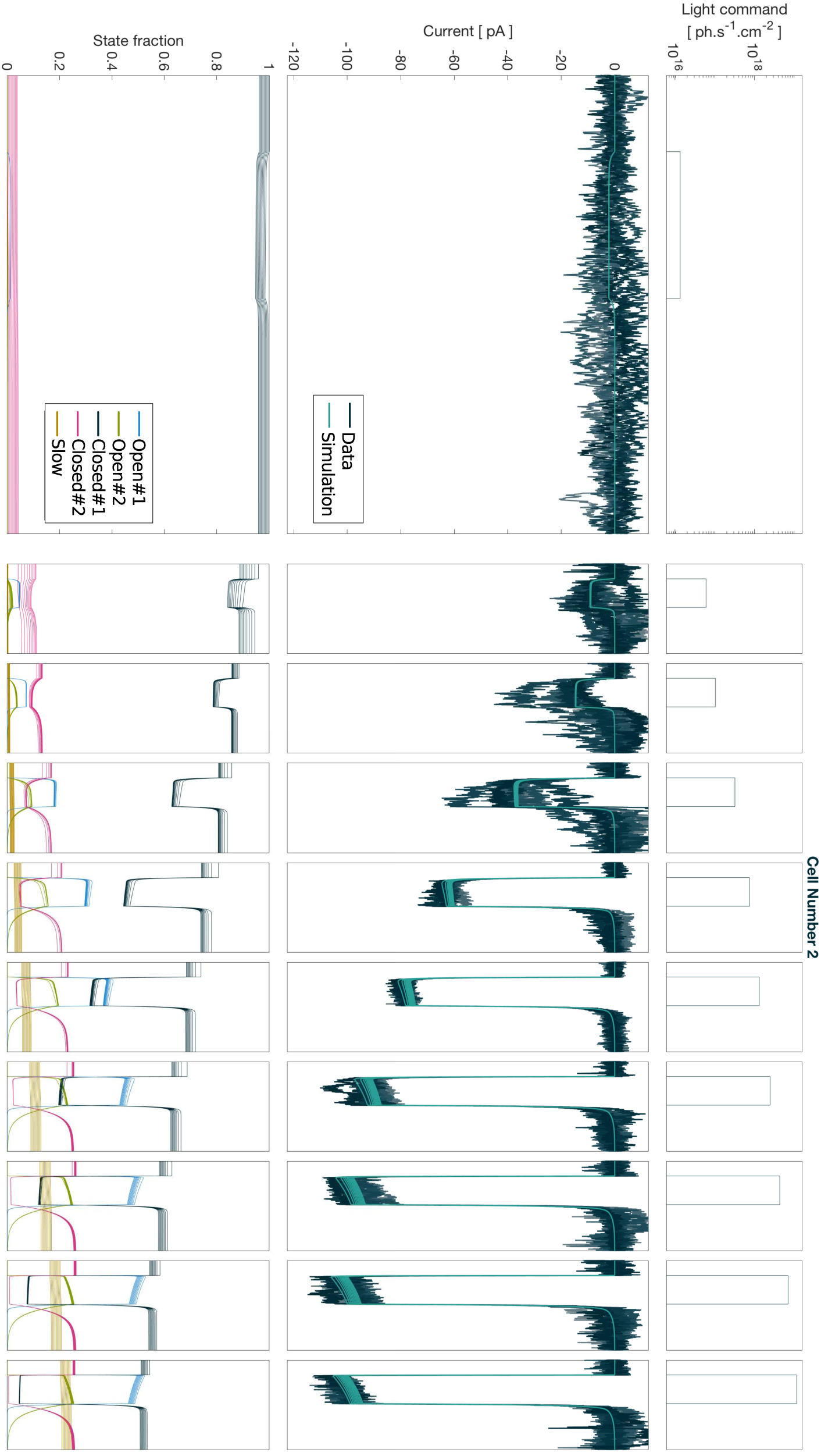
Data and simulation comparison Cell 2.

**Fig 10.**
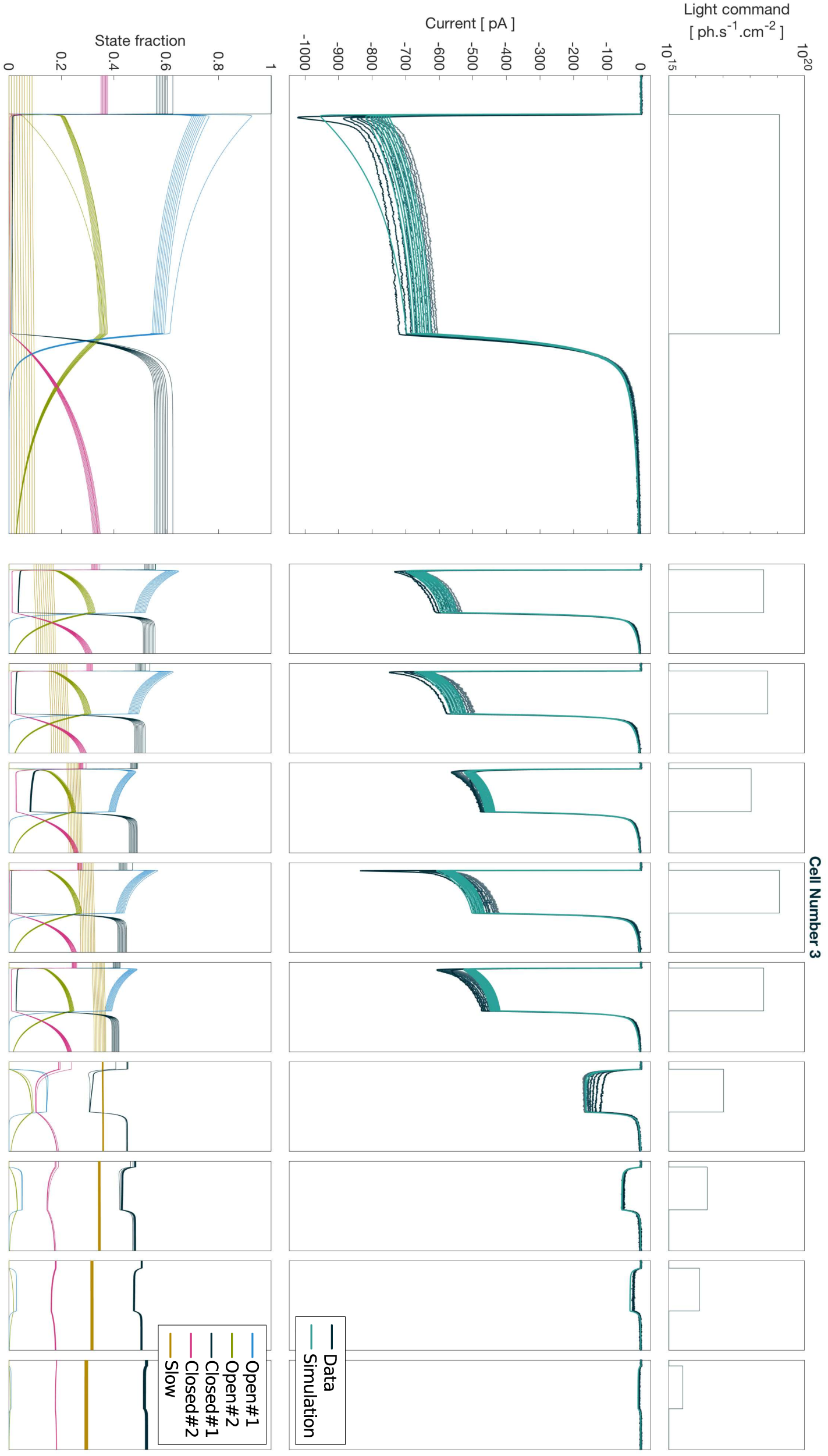
Data and simulation comparison Cell 3.

**Fig 11.**
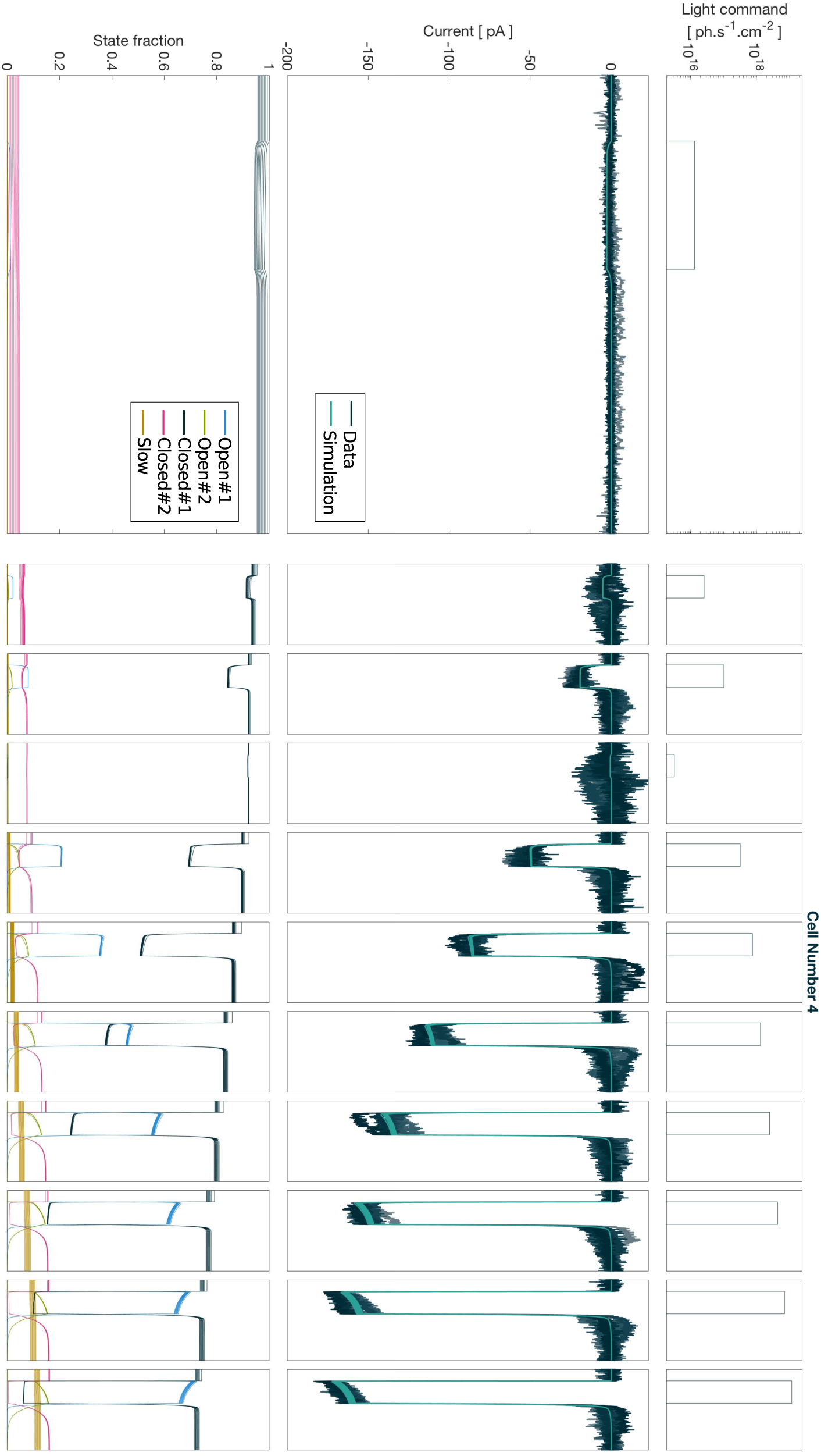
Data and simulation comparison Cell 4.

**Fig 12.**
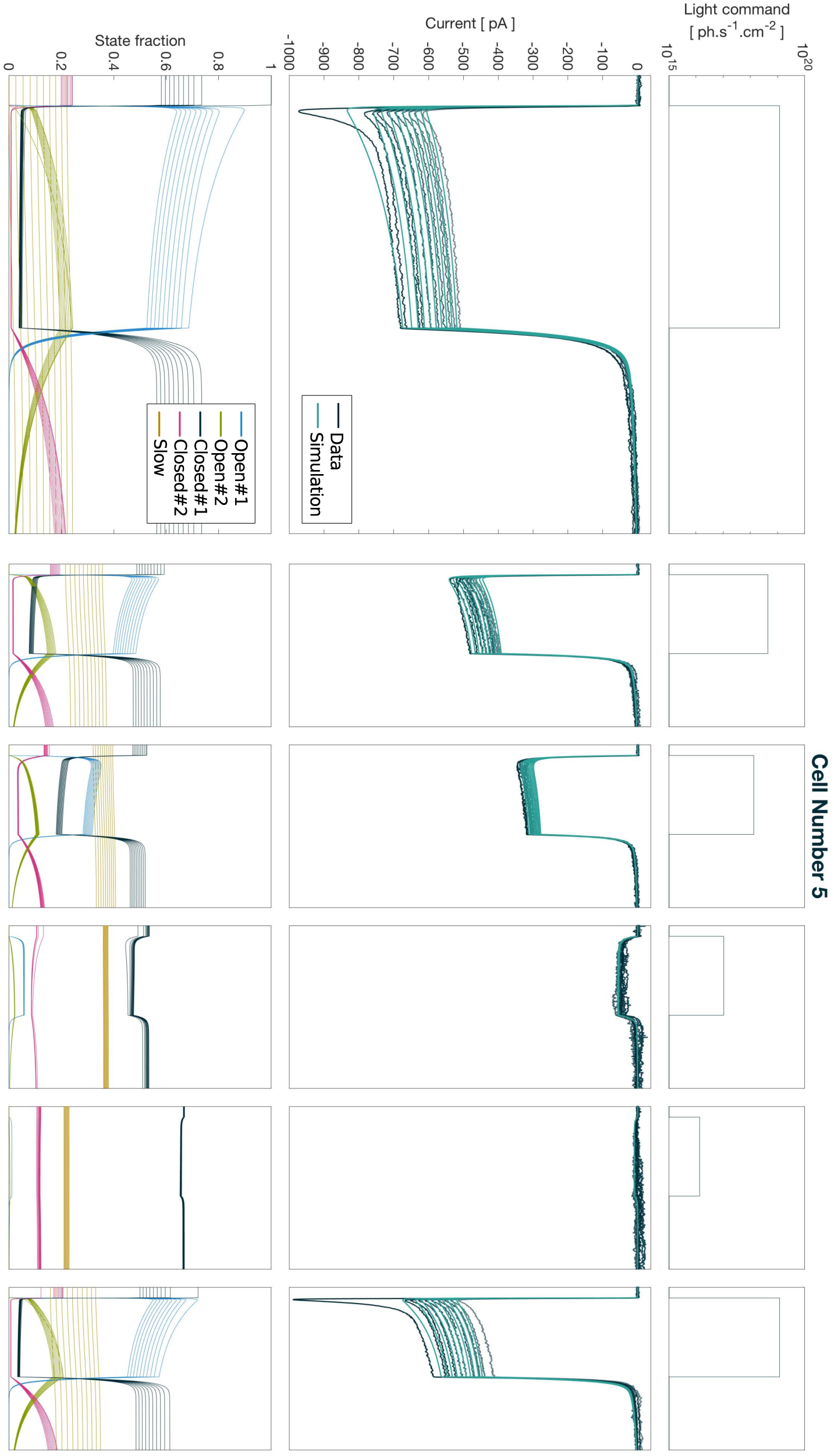
Data and simulation comparison Cell 5.

**Fig 13.**
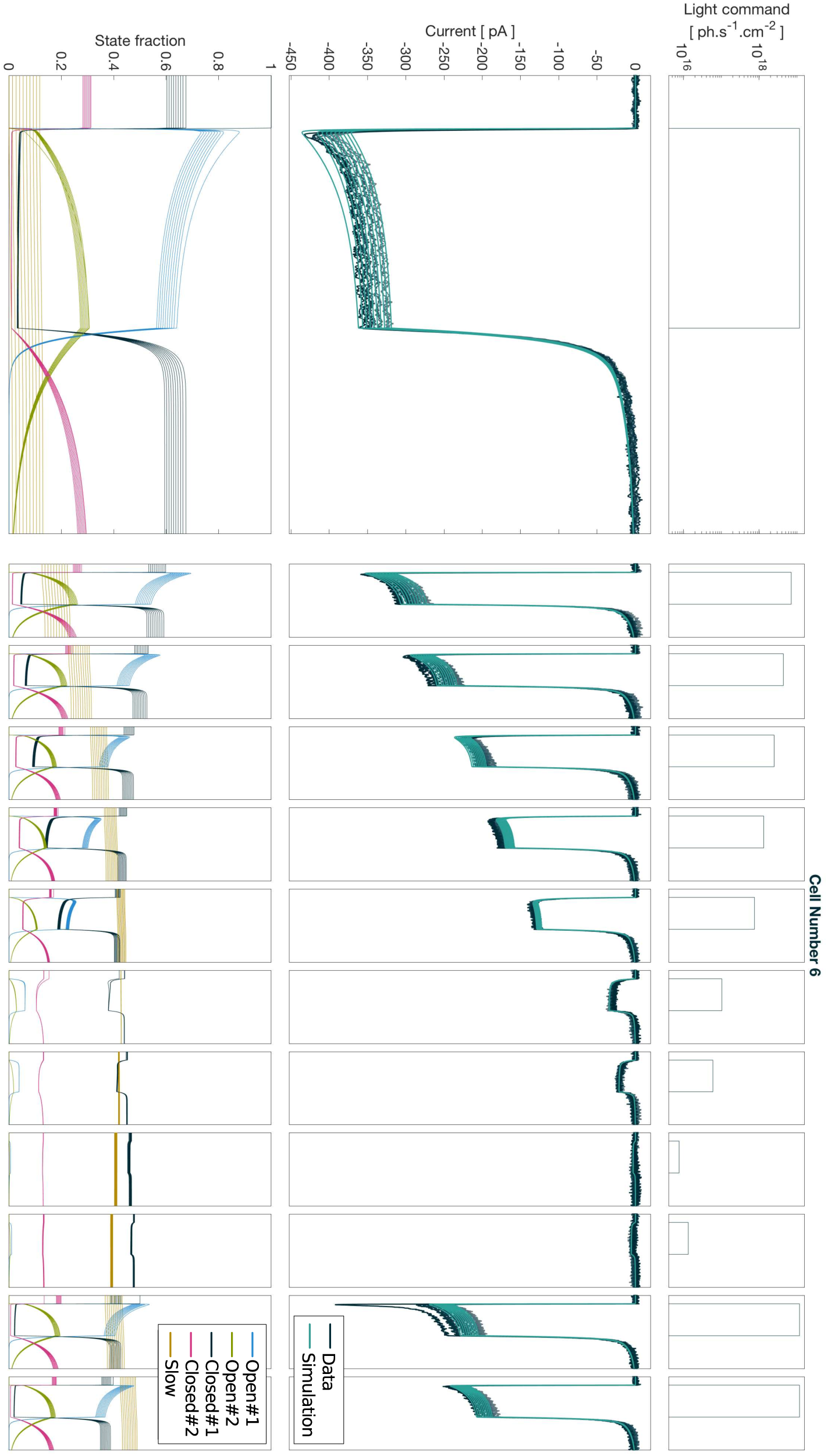
Data and simulation comparison Cell 6.

#### Activation curve

The five-state model allows the recovery of important information which is not directly available through simple statistical analysis of the raw data. One such information is the activation curve of the protein, i.e. the amplitude of the steady-state conductance of the protein as a function of light intensity.

Fig. (14) shows the activation curves for all parameters sets presented above. For each parameter set, two distinct activation curves are presented, the first one is the steady-state response at an intermediate timescale around 1 s for which the conductance has apparently reached a plateau, but which in fact slowly decays to a lower plateau due to the side reaction represented by the state S in Fig. (1, b). The second activation curve represents this real steady-state, including the side reaction.

The estimation of the slow time constant of the transition from the slow state S to the stable closed state C_1_ is quite imprecise because it is long relative to the time of the experiment and because of other phenomena involved in the experiment — clamping condition, physiological state of the cell, etc… — occur at the same timescale. However, assuming a sensible numerical value of 5 minutes for this time constant, we estimate that the long-term steady-state response is somewhere between 3 and 15 % of the middle-term value, the one that is observed when considering only short experiment.

#### ON time constants comparison

Contrary to the off dynamics, the preliminary studies of the on dynamics of the responses were not used to fit the parameters of the five-state model. As a safety check, we display side by side in Fig. (15) (i) the time constants estimated directly on the raw data as presented in paragraph and (ii) the time constants of the five sets of parameters introduced above.

**Fig 14.**
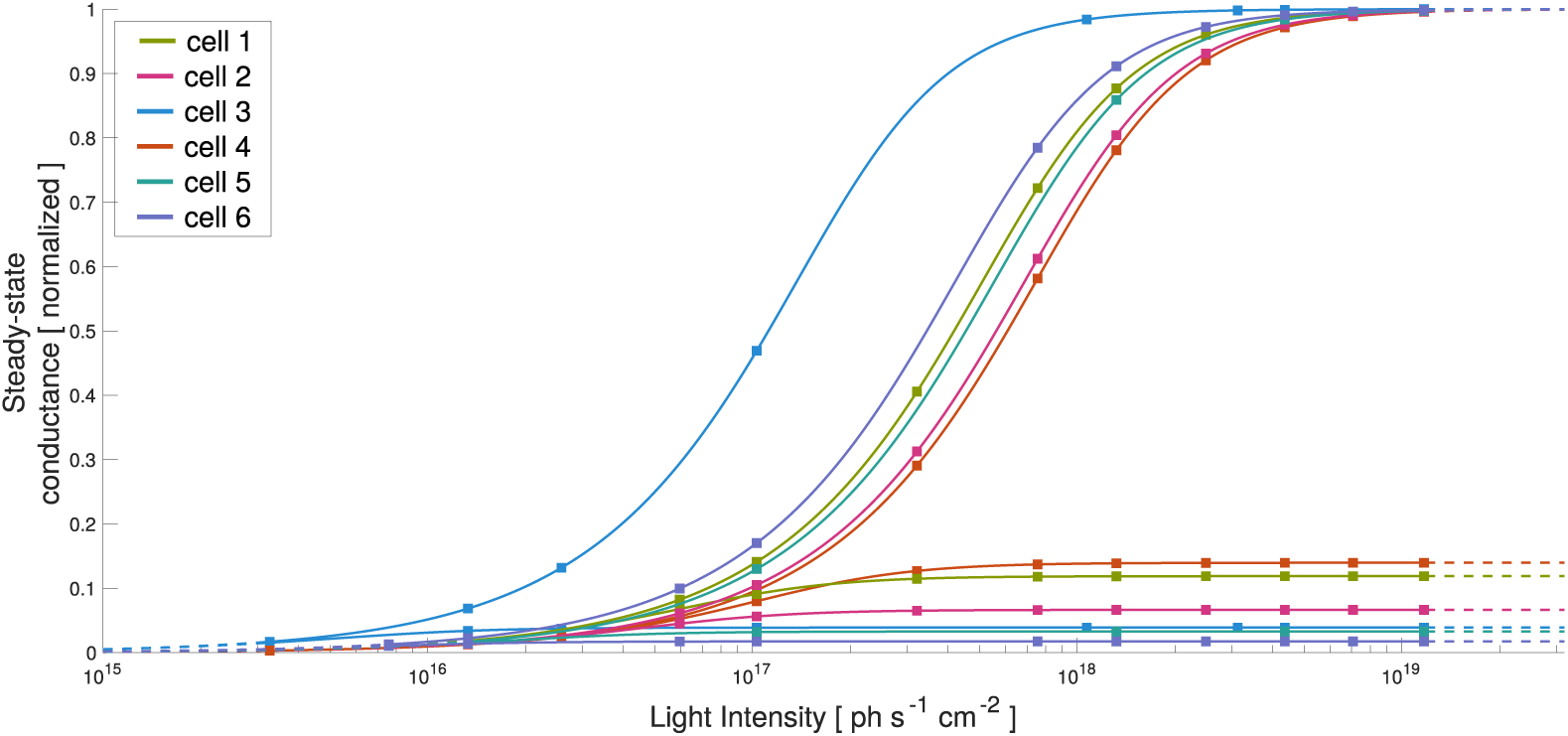
Activation curve of ChrimsonR at mid term steady-state. The curves are derived from the Five-state model fitted on data from four different cells, for which the response was observed on a wide range of light intensities. The curves were obtained theoretically according to the procedure described in the *Activation curve* paragraph and are normalized with respect to the limit value of medium-term activation curve. The dots on each curve represent the light intensity values which were present in the stimulation protocol on which the model was fitted. Thus, the curve is shown in dotted line outside the range of actual stimulation values. ChrimsonR starts to respond significantly around 10^16^ ph.s^−1^.cm^−2^ and saturates above 10^18^ ph.s^−1^.cm^−2^. Beyond this limit, the steady-state value of the conductance is unaffected. Only the dynamics are impacted.

**Fig 15.**
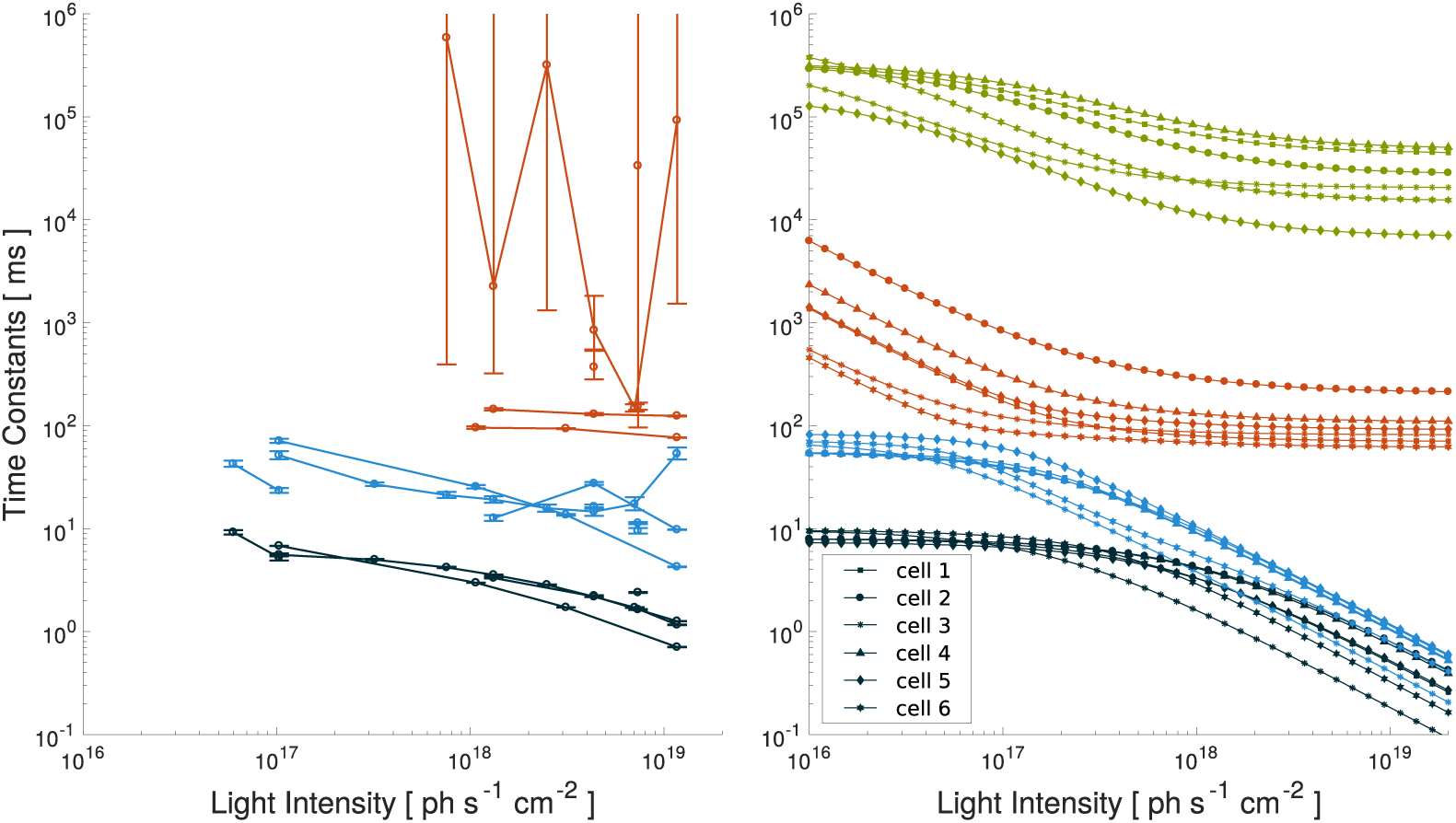
On dynamics comparison. Time constants of the linear combination of exponential functions underlying the responses to square pulses of light stimulation. (left) time constants are estimated directly on the data. Depending on the intensity of the light command, either two or three distinct time constants are estimated. (right) time constants derived from the five-state model dynamics. The time constants are computed as the non-zero eigenvalues of the differential operator governing the evolution of the probability distribution of the Markov chain. Note that this method reveals a fourth time constant. This fourth time constant represents the dynamics at which the open states equilibrate with the slow state S.

The figure shows that the dynamics are comparable between the recorded and simulated responses on the range of recorded light intensities (black, blue and orange curves). The time constants derived directly from the model are slightly quicker than the ones estimated on the data. Computing the time constants through the model allows us to reveal the dynamics at with the open states equilibrate with the slow state S (green curve). It represents the timescale at which we slide from the medium-term to the long-term steady-state (see Fig. 14).

## Discussion

In this paper, we propose a five-state Markov kinetic model describing the conductance response of channelrhodopsins ChrimsonR to an arbitrary light stimulus. The model was chosen to capture the main features of the experimental observations: activation curve, on and off kinetics, medium and long term light adaptation. These are the key features to design stimulation algorithms, i.e. setting the light stimulation signal in order to induce a given response. Our predictive model provides an essential tool for both medical and experimental applications.

The model was designed in the direct lineage of similar work, carried out over the past decade [11–13, 27], but also taking into account the more recent conclusion on the actual, more complex, photocycle of channelrhodopsins [25]. Additionally, our analysis is based on a set of voltage-clamp experiments on ChrimsonR-transfected HEK293 cells at a holding potential of-60 mV and at physiological temperature (for direct use in medical applications). The cells were subjected to rich stimuli, including a wide range of light intensities and temporal scales. These experiments allowed direct observation of the response amplitude and dynamics of the ion channels to light intensities covering the whole response range of the channels and dynamics over scales ranging from the millisecond to several minutes.

Direct observation of the data revealed that there is a fast light adaptation mechanism that has not been described previously. It was already known, from the shape of the conductance responses to light pulses, that the conductance does not reach a plateau directly, but rather peaks first before decaying to a stable value. This decay has been described as light adaptation and corresponds to an equilibration between the two cycles. We additionally report that the intensity of the light pulse influences the ratio between the two cycles, and in turn, the resulting ratio between the populations in the two closed states D470 and D480 after the light is turned off. Finally, this ratio determines the shape of the first part of the response to a new light pulse. This phenomenon results in a mechanism where channels adapted to high light intensities tend to respond with slower dynamics to new stimulation.

This feature is not captured by our five-state model, and we did not manage to capture it yet with other n-state models that we tested along the way. This underlines that the complex photocycle of channelrhodopsin is not fully understood and that work remains to be done before we establish the precise functional outline of the photocycle. However, we believe that once the photocycle is understood, fitting it to voltage-clamp data of the different known channelrhodopsin will to be quite straightforward. We also believe that numerical simulation is an essential tool in the search for the photocycle, confirming, invalidating or challenging hypotheses stemming from spectroscopic studies (which provides valuable information regarding the time constants of the existing reactions) and crystallography (which identifies the possible chemical reactions, changes of conformation and associate them with the structure of the protein).

Despite this limitation, our predictive model capture the key feature needed to design stimulation algorithms for both medical and experimental applications. Among these are the dynamics of the channels and the evolution of the on and off time constants with light intensity. This is critical in order to set the temporal frequency range on which to stimulate the photosensitive cells. The amplitude of the response and its relationship to light intensity is also fundamental since it directly influences the spiking mechanism in targeted neurons. However, the relationship between channel response amplitude and spike rate is not completely straightforward: it requires knowledge of the expression level of the channel population within the neuron and of the neuron’s spiking mechanism. This will be the subject of future work. Finally, our data unveils a slow adaptation mechanism. It leads to an attenuation of the response upon stimulation. The higher the stimulation, the higher the attenuation. Efficient stimulation algorithms therefore need to carefully manage the tradeoff between the efficiency of the response now and in the future. In this perspective, our model is a key tool in guiding the design of such algorithms and in assessing their efficiency.

## References

1. Boyden ES. Optogenetics and the future of neuroscience. Nature Neuroscience. 2015;18(9):1200–1201. doi:10.1038/nn.4094.

2. Klapoetke NC, Murata Y, Kim SS, Pulver SR, Birdsey-Benson A, Cho YK, et al. Independent optical excitation of distinct neural populations. Nature Methods. 2014;11(3):338–346. doi:10.1038/nmeth.2836.

3. GenSight Biologics;. Available from: https://www.gensight-biologics.com.

4. Destexhe A, Mainen ZF, Sejnowski TJ. Synthesis of models for excitable membranes, synaptic transmission and neuromodulation using a common kinetic formalism. Journal of computational neuroscience. 1994;1(3):195–230. doi:10.1007/BF00961734.

5. Müller M, Bamann C, Bamberg E, Kühlbrandt W. Projection structure of channelrhodopsin-2 at 6 °A resolution by electron crystallography. Journal of Molecular Biology. 2011;414(1):86–95. doi:10.1016/j.jmb.2011.09.049.

6. Kato HE, Zhang F, Yizhar O, Ramakrishnan C, Nishizawa T, Hirata K, et al. Crystal structure of the channelrhodopsin light-gated cation channel. Nature. 2012;482:369–374. doi:10.1038/nature10870.

7. Hososhima S, Sakai S, Ishizuka T, Yawo H. Kinetic evaluation of photosensitivity in Bi-stable variants of chimeric channelrhodopsins. PLoS ONE. 2015;10(3):1–14. doi:10.1371/journal.pone.0119558.

8. Krause BS, Grimm C, Kaufmann JCD, Schneider F, Sakmar TP, Bartl FJ, et al. Complex Photochemistry within the Green-Absorbing Channelrhodopsin ReaChR. Biophysical Journal. 2017;112(6):1166–1175. doi:10.1016/j.bpj.2017.02.001.

9. Schneider F, Gradmann D, Hegemann P. Ion selectivity and competition in channelrhodopsins. Biophysical Journal. 2013;105(1):91–100. doi:10.1016/j.bpj.2013.05.042.

10. Lórenz-Fonfría VA, Heberle J. Channelrhodopsin unchained: Structure and mechanism of a light-gated cation channel. Biochimica et Biophysica Acta-Bioenergetics. 2014;1837(5):626–642. doi:10.1016/j.bbabio.2013.10.014.

11. Hegemann P, Ehlenbeck S, Gradmann D. Multiple photocycles of channelrhodopsin. Biophysical journal. 2005;89(6):3911–8. doi:10.1529/biophysj.105.069716.

12. Nikolic K, Grossman N, Grubb MS, Burrone J, Toumazou C, Degenaar P. Photocycles of channelrhodopsin-2. Photochemistry and Photobiology. 2009;85(1):400–411. doi:10.1111/j.1751-1097.2008.00460.x.

13. Evans BD, Jarvis S, Schultz SR, Nikolic K. PyRhO: A Multiscale Optogenetics Simulation Platform. Frontiers in Neuroinformatics. 2016;10(March):1–19. doi:10.3389/fninf.2016.00008.

14. Lin B, Masland RH, Strettoi E. Remodeling of cone photoreceptor cells after rod degeneration in rd mice. Experimental eye research. 2009;88(3):589–599. doi:10.1016/j.exer.2008.11.022.

15. Bamann C, Gueta R, Kleinlogel S, Nagel G, Bamberg E. Structural guidance of the photocycle of channelrhodopsin-2 by an interhelical hydrogen bond. Biochemistry. 2010;49(2):267–278. doi:10.1021/bi901634p.

16. Kuhne J, Eisenhauer K, Ritter E, Hegemann P, Gerwert K, Bartl F. Early Formation of the Ion-Conducting Pore in Channelrhodopsin-2. Angewandte Chemie. 2015;54:4953–4957. doi:10.1002/anie.201410180.

17. Gunaydin LA, Yizhar O, Berndt A, Sohal VS, Deisseroth K, Hegemann P. Ultrafast optogenetic control. Nature Neuroscience. 2010;13(3):387–392. doi:10.1038/nn.2495.

18. Watanabe HC, Welke K, Sindhikara DJ, Hegemann P, Elstner M. Towards an understanding of channelrhodopsin function: Simulations lead to novel insights of the channel mechanism. Journal of Molecular Biology. 2013;425(10):1795–1814. doi:10.1016/j.jmb.2013.01.033.

19. Bamann C, Kirsch T, Nagel G, Bamberg E. Spectral Characteristics of the Photocycle of Channelrhodopsin-2 and Its Implication for Channel Function. Journal of Molecular Biology. 2008;375(3):686–694. doi:10.1016/j.jmb.2007.10.072.

20. Radu I, Bamann C, Nack M, Nagel G, Bamberg E, Heberle J. Conformational Changes of Channelrhodopsin-2. The Journal of the American Chemical Society. 2009;446(7136):7313–7319.

21. Lorenz-Fonfria VA, Resler T, Krause N, Nack M, Gossing M, Fischer von Mollard G, et al. Transient protonation changes in channelrhodopsin-2 and their relevance to channel gating. Proceedings of the National Academy of Sciences. 2013;110(14):E1273–E1281. doi:10.1073/pnas.1219502110.

22. Berndt A, Prigge M, Gradmann D, Hegemann P. Two open states with progressive proton selectivities in the branched channelrhodopsin-2 photocycle. Biophysical Journal. 2010;98(5):753–761. doi:10.1016/j.bpj.2009.10.052.

23. Stehfest K, Hegemann P. Evolution of the Channelrhodopsin Photocycle Model. ChemPhysChem. 2010; p. 1120–1126. doi:10.1002/cphc.200900980.

24. Ritter E, Piwowarski P, Hegemann P, Bartl FJ. Light-dark Adaptation of Channelrhodopsin C128T Mutant. The Journal of Biological Chemistry. 2013;288(15):10451–10458. doi:10.1074/jbc.M112.446427.

25. Bruun S, Stoeppler D, Keidel A, Kuhlmann U, Luck M, Diehl A, et al. Light Dark Adaptation of Channelrhodopsin Involves Photoconversion between the all-trans and 13-cis Retinal Isomers. Biochemistry. 2015;doi:10.1021/acs.biochem.5b00597.

26. Dalkara D, Byrne LC, Klimczak RR, Visel M, Yin L, Merigan WH, et al. In Vivo-Directed Evolution of a New Adeno-Associated Virus for Therapeutic Outer Retinal Gene Delivery from the Vitreous. Science Translational Medicine. 2013;5(189):189ra76. doi:10.1126/scitranslmed.3005708.

27. Grossman N, Nikolic K, Toumazou C, Degenaar P. Modeling study of the light stimulation of a neuron cell with channelrhodopsin-2 mutants. IEEE Transactions on Biomedical Engineering. 2011;58(6):1742–1751. doi:10.1109/TBME.2011.2114883.

